# The sense of agency in near and far space: where do we stand?

**DOI:** 10.64898/2025.12.22.695728

**Authors:** Gaiqing Kong, Marine Vernet, Alessandro Farnè

## Abstract

Modern technology frequently places the consequences of our actions at a distance (e.g., remote surgery, smart-home control, virtual reality). Does spatial distance between an action and its outcome weaken the sense of agency (SoA) - the feeling of control over one’s actions and consequences? Two recent studies answered “yes”, reporting stronger temporal binding (TB) in near than far space and interpreting this as greater implicit agency (Jenkins & Obhi, 2021; Mariano et al., 2024). A third study - our own work with a similar paradigm - found no distance effect (Kong et al., 2024). Here we (i) provide a rigorous side-by-side methodological comparison of the three studies, (ii) argue why a direct test to establish a distance modulation of TB (the Near – Far difference of the Active – Passive delta) should be performed in order to reach meaningful conclusions, and (iii) report new re-analyses of our data and direct tests on the two target studies. Overall, current evidence does not support a distance effect on SoA. Our reassessment provides alternative explanations that converge with available evidence suggesting that distance may influence temporal interval perception, but that effect is independent of action intention and therefore of agency.

**Public Significance Statement:** In our increasingly connected world, we often interact with devices and influence events that are physically distant from us - like controlling smart appliances remotely or engaging in virtual reality experiences. Does the distance between us and the effects of our actions change how much we feel in control of the outcomes of our actions? Recent studies have suggested that we feel less responsible when action outcomes appear in far space compared to near space. However, methodological limitations in these studies, as well as our own results with a similar paradigm, challenge the validity of this claim. We aim to contribute to the crucial debate on the role of spatial distance on humans’ feeling of responsibility by providing a respectful though critical analysis of recent findings and offering recommendations for future research.

## Introduction

Modern technology has dramatically transformed how we interact with the world, enabling actions performed in near space to produce far-reaching consequences. Despite this shift, the role of spatial distance - specifically where the outcomes of our actions are perceived - has been largely overlooked in research on the sense of agency (SoA). SoA is a fundamental component of self-consciousness, allowing individuals to recognise themselves as the agents of their actions and their ensuing consequences (Haggard, 2017; Haggard & Chambon, 2012). Understanding how spatial context influences SoA is particularly relevant in today’s increasingly digital and remote environments, where the boundaries between near and far space are becoming increasingly blurred. However, this question has remained largely underexplored in previous research.

Mariano et al. (2024) sought to address this question by manipulating the spatial context in which participants perceived their action outcomes, using both illusory computer-based and realistic settings. In their study, participants performed an active or passive button press to light up a lightbulb. The authors used an implicit measure of agency, the “intentional binding”, which refers to the phenomenon where individuals perceive the temporal interval between their voluntary action and its consequence as shorter than the interval between two events in which they are not voluntarily involved. The authors reported that such binding was greater when outcomes occurred in near space compared to far space, regardless of the ecological validity of the setting. In contrast, explicit agency judgments were not significantly affected by spatial distance. The authors concluded that spatial proximity - whether realistic or illusory - is essential for feeling implicitly responsible for the consequences of our actions.

Previously, Jenkins and Obhi (2021) also explored the relationship between perceived action-outcome distance and the sense of agency using the temporal binding effect, a more general term encompassing intentional binding (IB). In their online study, they manipulated the perceived near and far distances of action outcomes through perspective cues. Participants triggered the appearance of a circle at different perceived distances by a voluntary keypress or observed the same circle being triggered by a computer-generated tone. Jenkins and Obhi found that participants’ interval estimates - the perceived time between action and outcome - increased with the perceived distance of the outcome. However, this increase was not restricted to conditions involving active movement within a 3D perspective, it was also observed in other conditions. The authors carefully stated that their results point to a link between intentionality and distance, in the direction of a reduced sense of agency for outcomes that appear further away, while acknowledging the limitations of their study and the need for further research.

Having been researching on this topic without finding evidence for an effect of spatial proximity on “intentional binding” (Kong et al., 2024), we closely inspected and provided a direct reanalysis of the available work, including our own. Specifically, we analytically assess the work of both Mariano et al. (2024) and Jenkins and Obhi (2021), focusing on data analyses and the conclusions drawn, and we also compare their results with our own previous data (Kong et al., 2024), which did not support the effect of distance on temporal binding in contexts involving intentional movements. We end by suggesting directions for future research in this domain.

### How to measure temporal binding and assess its modulation by spatial proximity

Both Jenkins and Obhi (2021) and Mariano et al. (2024) investigated the effect of spatial distance on the SoA by manipulating the perceived location of action outcomes. Jenkins and Obhi manipulated spatial distance using visual perspective cues in an online study, presenting outcomes that appeared near or far from the participant’s viewpoint (Table 1). The spatial proximity of the outcome was cued using a 3D perspective (simulating near vs. far space by varying the relative size and position of the visual elements). As a control, a flat 2D condition without perspective cues was used. Mariano et al. used both computer-based environments and realistic settings to present action outcomes in near (∼35 cm) and far (∼70 cm) space (Table 1). Both studies employed the temporal binding paradigm with the interval estimation procedure. In this paradigm, the perceived time interval between an action and its outcome is typically shortened for voluntary (active) movements compared to involuntary (passive) movements, or baseline conditions, and this temporal compression is taken as an implicit measure of SoA (Moore & Obhi, 2012). Jenkins and Obhi (2021) employed intervals of 200, 500, and 800 ms, while Mariano et al. (2024) used intervals of 200, 400, and 600 ms.

**Table 1.** Three studies have brought TB to bear on spatial distance.

| Study | Display & distance cue | First-event type | Delays (ms) | Core claim |
| --- | --- | --- | --- | --- |
| **Jenkins & Obhi (2021)** | PC monitor; linear-perspective cues; *flat* "no-perspective" control | Active keypress; External event | 200 / 500 / 800 | 200 ms intervals judged shorter when the circle appears "near" |
| **Mariano et al. (2024)** | (a) PC, retinal-size cue (35 cm vs 70 cm) and (b) Real table-top | Active keypress, Passive keypress | 200 / 400 / 600 | Larger TB at 200 ms in near vs far space |
| **Kong et al. (2024)** | Immersive VR; binocular disparity (45 cm vs 4.5 m) | Active keypress, Passive keypress, External event | 150 / 450 / 750 | Distance does not modulate TB |

As a premise that we have addressed in previous work, we wish to note that, despite still being widely used, the intentional binding phenomenon is increasingly recognized as multifaceted and potentially confounded. Recent studies have shown that intentional binding may arise from factors such as causality, predictability and somatosensory integration, rather than uniquely reflect intentionality and the voluntariness of action per se (Buehner, 2015; Buehner & Humphreys, 2009; Dewey, 2024; Gutzeit et al., 2023; Kirsch et al., 2019; Kong et al., 2024; Suzuki et al., 2019). Consequently, researchers should be cautious when using “intentional binding” as a reliable proxy for the sense of agency. Both Mariano et al. (2024) and Jenkins and Obhi (2021) acknowledge these criticisms. However, Mariano et al. (2024) argue that there are no alternative explanations, and state “the measure is achieved via the comparison of active intentional versus passive movements and it is systematically larger for active movements (i.e., shorter perceived temporal interval between actions and consequences)” (p. 2). Jenkins and Obhi (2021) note alternative interpretations that question whether intentional binding necessarily reflects agency, such as causal inference and multisensory integration. Yet, they also suggest that these criticisms are not definitive and that there remains evidence supporting temporal binding as a measure influenced by intentional actions. We concur that comparing a voluntary movement condition with an involuntary baseline condition is critical for measuring temporal binding (Haggard et al., 2002; Moore & Obhi, 2012), though it may not alone justify labelling the effect as ‘intentional’. The importance of relying on the comparison between an active condition and a baseline forms the basis of our assessment of the two previous studies (Jenkins & Obhi, 2021; Mariano et al., 2024).

Mariano et al. concluded that the intentional binding effect is larger in near space compared to far space, based on a pairwise comparison of active trials with a 200 ms action-outcome delay between near and far space (t (54) = -3.57, Bonferroni-corrected p = 0.05, Cohen’s d = -0.95). While passive trials were included in their ANOVA, the conclusion was drawn from only comparing active trials between distances and passive trials between distances separately, which we deem problematic. Intentional binding is defined as the perceived shortening of the time interval between an action and its outcome when the action is intentional, compared to unintentional actions, or mere action observation. Therefore, measuring the intentional binding effect requires calculating the *difference* in temporal interval estimates between active and passive trials. When investigating the influence of the spatial distance of an action outcome on the SoA, it is essential to access a difference-of-differences (i.e., a delta measure):

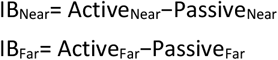

A spatial modulation of binding therefore requires a statistical comparison of IB_Near_ versus IB_Far_, rather than comparing active-only and/or passive-only estimates across distances.

Even when a significant Action × Distance interaction is observed in an ANOVA, this alone does not justify concluding a distance effect on binding. Such a conclusion requires specific post-hoc tests confirming that the difference between active and passive trials is significantly larger in near space compared to far space (Fig. 1). Indeed, the values reported in Mariano et al. indicate that the actual difference in intentional binding (*Δ*IB) between near and far space is modest - approximately 11 ms in the computer-based condition and only ∼1 ms in the realistic setting. Crucially, this difference was not directly tested: the well-established method for measuring intentional binding, where each condition must be assessed relative to its corresponding baseline (Haggard et al., 2002), was not applied.

**Figure 1.**
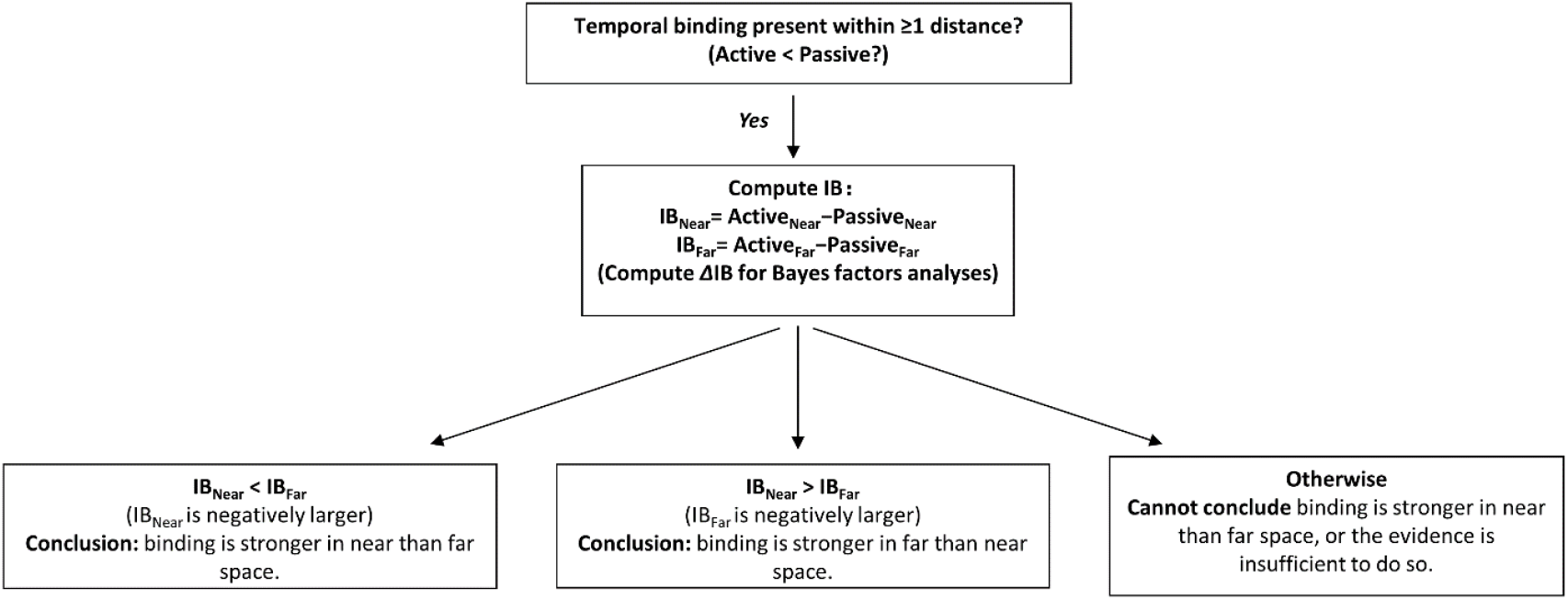
Decision tree for testing whether spatial distance modulates the temporal binding effect. First, establish a binding effect within at least one distance. Then compute the binding effects for Near and Far, respectively, and statistically compare IB_Near_ and IB_Far_ using a null-hypothesis significance testing (NHST) approach. For Bayesian inference, assess the modulation directly via the ΔIB (difference-of-differences) contrast. We would like to note: ‘significant in Near, non-significant in Far’ is not evidence for a modulation; only a supported non-zero *Δ*IB warrants the conclusion (Gelman & Stern, 2006; Pálfi & Dienes, 2020).

Taking this opportunity, we also wish to clarify the repeated use and the definition of the term “time compression” in some studies (Mariano et al., 2024; Zapparoli, Seghezzi, Devoto, et al., 2020; Zapparoli, Seghezzi, Zirone, et al., 2020) to avoid misunderstanding. Defining “Time compression (TC)” as the raw difference between the estimated and objective physical interval does not correspond to temporal binding (see Box 1). Such a quantity is more accurately described as judgement error or a perceptual shift (Haggard et al., 2002; Kong et al., 2017; Kühn et al., 2013). Again, intentional binding requires comparing active versus appropriate baseline conditions; absolute estimation error is not equivalent to intentional binding. It is also worth noting another misinterpretation, particularly common in the intentional binding literature, namely, that intentional binding measures the implicit sense of agency (see Box 1). This distinction is worth noting, as it appears increasingly frequently in the literature (Mariano et al., 2024; Zapparoli, Seghezzi, Devoto, et al., 2020; Zapparoli, Seghezzi, Zirone, et al., 2020).

#### Box 1.

**Two common misinterpretation in the intentional binding literature**

Misinterpretation 1: Difference between “Temporal compression” and “Temporal binding” Some studies define *temporal compression* (TC) as the raw difference between the actual duration and the estimated duration of the action-outcome delay, and equate this quantity with intentional binding. However, this quantity reflects *estimation/judgement error* or *perceptual shift/bias*, not intentional binding. Intentional binding is the reduction in perceived interval for voluntary actions relative to an appropriate baseline. Therefore, this absolute estimation error ≠ intentional binding. Humans systematically misjudge intervals (longer or shorter) in many contexts.

Misinterpretation 2: Difference between “implicit measure of the sense of agency” and “measure of the implicit sense of agency”

It is worth noting a common misconception, particularly widespread in the intentional binding literature, namely, that intentional binding measures the implicit sense of agency. The measure is often described as “implicit” because it does not involve explicit reflection on agency; however, this does not necessarily mean that it measures an implicit agency.

Jenkins and Obhi (2021) also reported a distance effect, finding that the 200 ms interval was estimated to be shorter in near space compared to far space. However, this distance effect occurred irrespective of the presence or absence of intentional movement and was not directly linked to the temporal binding effect they observed at the shortest 200 ms delay. When all delays were combined in the 3D condition, intervals were estimated to be shorter in near space than far space for the active movement task but not for the baseline task. However, no temporal binding was detected in the first place when comparing the active movement to the baseline tasks when all delays were combined in the 3D condition. Furthermore, the distance effect was also present in other conditions, such as the flat, no-perspective condition. The authors acknowledged these limitations and raised caution when suggesting that agency is reduced for outcomes that appear further away. Indeed, this conclusion requires further evidence.

In summary, we argue that any claim that spatial distance modulates intentional binding - and by extension, the sense of agency - requires the following minimal criteria (**Fig. 1**): (1) Presence of a temporal binding effect: Active < Passive within at least one spatial condition; (2) A specific comparison of binding magnitude across spatial conditions, i.e., (Active_Near_−Passive_Near_) differs from (Active_Far_−Passive_Far_) (Gelman & Stern, 2006; Palfi & Dienes, 2020). Previous work claiming a distance effect on SoA (Jenkins & Obhi,2021; Mariano et al., 2024) did not meet all of these criteria.

Finally, the issue is not only methodological but also theoretical: the passive condition serves as a baseline and its subtraction ensures that any observed effects would reflect changes in binding rather than general differences in time perception. We contend that the distance effect reported may reflect differences in temporal interval perception between the near and far conditions, rather than a spatial modulation of temporal binding. Below, reanalyses of previously published datasets directly assess the evidence for each of these competing hypotheses.

### Direct test: does spatial distance really modulate temporal binding?

Similar to previous work, in a recent study (Kong et al., 2024) we employed an immersive virtual reality (VR) environment to manipulate the location of a light-bulb, presenting action outcomes (the bulb lightening) at 45 cm (near) and 4.5 m (far) (Table 1, **Fig. 2**). The three delayed intervals tested were 150, 450, and 750 ms. In an attempt to isolate the role potentially played by ‘intentionality’ in the temporal binding paradigm, we introduced a cue to give a similar level of predictability to the first event (Kirsch et al., 2019; Kong et al., 2024; Saad et al., 2024), which could be an active keypress, a passive keypress, or an external sensory event (e.g., an off-bulb). More precisely, in the active movement task, participants performed a voluntary key press which switched on a light bulb. In the passive movement task, the experimenter pressed the participant’s left index finger down onto the button to switch on the light bulb. In the external sensory event task, participants merely observed the appearance of a light bulb which turned on after a delay. Using Bayes factor (BF) analyses, we found no evidence of intentional binding at any of these intervals when comparing active and passive movement tasks. Most notably, robust temporal binding emerged when comparing both active and passive movement tasks to an externally triggered baseline task. Because these results did not differ based on whether the action outcome occurred in near or far space - a question of great interest to us - we pooled the data from both distances in our published article.

**Figure 2.**
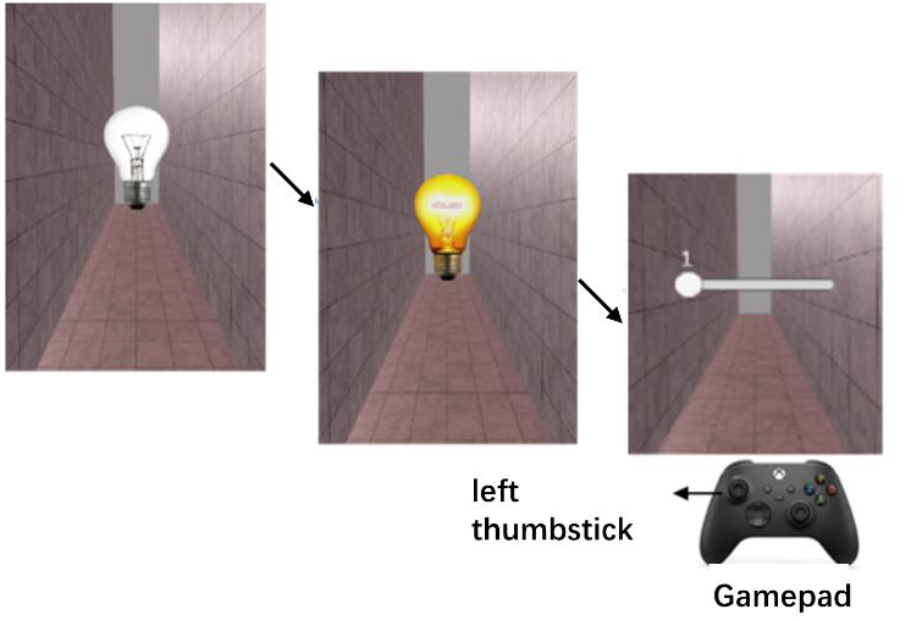
Experimental Setup in Our Study. In our experiment (Kong et al., 2024), participants viewed a virtual scene through a head-mounted display (Oculus Rift CV1). The action outcome was a visual effect (the lighting of a light bulb) which was presented at two different distances: near (45 cm) and far (4.5 m). During each trial, an unlit bulb was turned on by different first events depending on the task: an active or passive keypress, or an external sensory event (an off-bulb). In the active movement task, participants performed a voluntary keypress with their left index finger whenever they wanted. In the passive movement task, the experimenter pressed the participant’s left index finger down onto the button. The keypress and interval estimates were performed using a gamepad. Participants indicated their estimates by moving a visually depicted slider with the left thumbstick of the gamepad. In both active and passive movement tasks, the light bulb was illuminated when an active or passive keypress (button “A” on the gamepad) was made. After the lit bulb had appeared for one second, participants were asked to estimate the interval between the keypress and the onset of the bulb lighting. The three possible intervals were randomly set to 150 ms, 450 ms, or 750 ms. The slider ranged from a minimum value of 1 ms to a maximum of 1,000 ms, with the current value displayed in real-time as participants adjusted it. The near and far bulbs were displayed with retinal correction.

In light of Jenkins and Obhi (2021) and Mariano et al. (2024), we re-analyzed our previous data and conducted additional Bayes factors (BF) analyses to assess whether distance could still modulate “non-intentional” binding and temporal compression effects (**Fig. 3**). First, we report the results that concern temporal binding, analysed separately for the near and far distances, using the same assumptions outlined in Kong et al. (2024). Specifically, when comparing the active and passive movement tasks, the BF indicated substantial evidence for the null hypothesis at both distances, i.e. supporting the absence of intentional binding (Near: BF_HN(0,122.5)_ = 0.11, RR_B<1/3_ = [40, >3000]; Far: BF_HN(0,122.5)_ = 0.27, RR_B<1/3_ = [100, >3000]). However, we observed a robust temporal binding effect when comparing both active and passive movement tasks to an externally triggered sensory event at both distances (**Fig. 3**). For the active movement versus external sensory event task, the BF indicated substantial evidence for a difference, at both distances (Near: BF_HN(0,122.5)_ = 4.01, RR_B>3_ = [9, 167]; Far: BF_HN(0,122.5)_ = 3.29, RR_B>3_ = [10, 135]). Similarly, for the passive movement versus external sensory event task, the BF showed strong evidence for a difference, at both distances (Near: BF_U(13.5,122.5)_ = 24.32, RR_B>3_ = [1, 882]; Far: BF_U(13.5,122.5)_ = 4090, RR_B>3_ = [1, >3000]).

**Figure 3.**
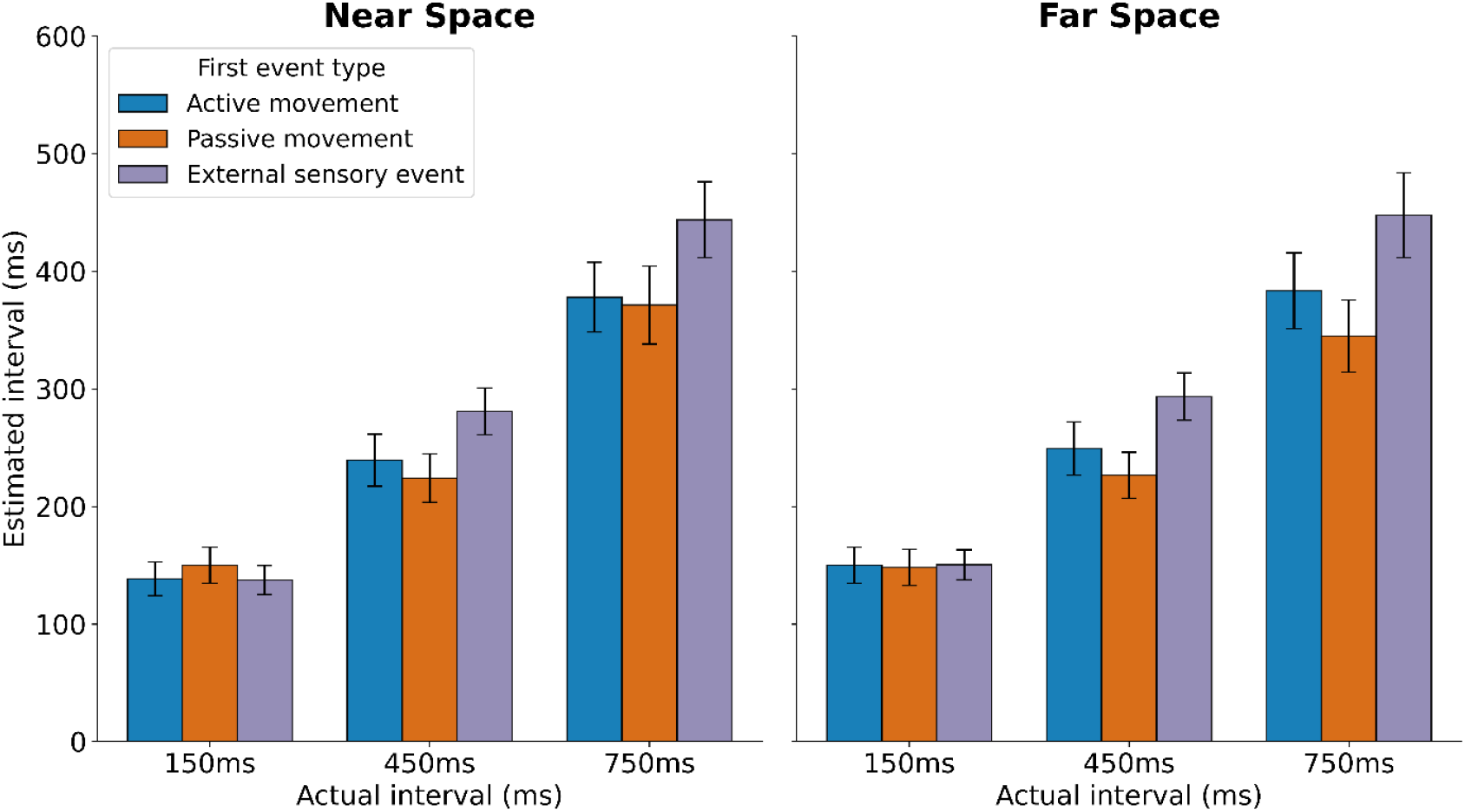
Estimated time intervals across near and far space for active movement, passive movement, and external sensory event tasks. The left panel represents results for the near space condition (45 cm), and the right panel represents the far space condition (4.5 m). The x-axis indicates the actual interval (150 ms, 450 ms, 750 ms), while the y-axis shows the estimated interval (ms). Data are color-coded to distinguish between the first event types: blue for active trials, orange for passive trials, and purple for external sensory events. Error bars indicate the standard error of the mean (SEM).

To examine whether non-intentional temporal binding was modulated by distance, we calculated the BF by using a half-normal distribution with a standard deviation of 17.265 ms, which was the size of the distance effect on temporal interval estimation in the active task, extracted from Figure 2 of Mariano et al. (2024). Using this prior, we found inconclusive evidence that spatial distance modulates the temporal binding between active movement and external event tasks (BF_HN(0,17.265)_ = 0.43, RR_1/3<B<3_ = [1, 22]). When testing each interval separately, the evidence was also all inconclusive for all delays (150ms: BF_HN(0, 17.265)_ = 0.50, RR_1/3<B<3_ = [1, 27]; 450ms: BF_HN(0, 17.265)_ = 0.69, RR_1/3<B<3_ = [1, 43]; 750ms: BF_HN(0, 17.265)_ = 0.74, RR_1/3<B<3_ = [1, 50]).

In contrast, there was substantial evidence that spatial distance modulated the temporal binding between passive movement and external event tasks (BF_HN(0,17.265)_ = 17.74, RR_B>3_ = [4, 178]), with temporal binding being smaller in near space than in far space – thus opposite to the expected distance effect. When testing each delay separately, there was also substantial evidence supporting the distance modulation in the opposite direction for the 150 ms delay (BF_HN(0, 17.265)_ = 5.896, RR_B>3_ = [5, 47]). For the 450 ms and 750 ms delays, the evidence was inconclusive (450ms: BF_HN(0, 17.265)_ = 1.26, RR_1/3<B<3_ = [1, 86]; 750ms: BF_HN(0, 17.265)_ = 2.35, RR_1/3<B<3_ = [1, 398]).^1^

However, a modulation analysis presupposes that the effect to be modulated is present. In our VR study (Kong et al., 2024), we did not observe intentional binding in the strict sense (Active − Passive), and therefore, those data cannot adjudicate whether intentional binding changes with distance.

In order to directly test whether spatial distance modulates temporal binding in the target literature, we computed the decisive difference-of-differences contrast *Δ*IB = (Active−Passive)_Near_−(Active−Passive)_Far_ at the critical 200-ms delay for both previous studies. For Mariano et al. (2024), raw participant data were not accessible. Using their Figure 2, we obtained ΔIB = −6.18 ms, and we estimated SE_Δ_ = 8.24 ms from their Supplementary data^2^. Using a half-normal distribution with a standard deviation of 17.265 ms (the size of the distance effect on temporal interval estimation in the active task extracted from their Figure 2), we found inconclusive evidence for a stronger binding effect in near space compared to far space (BF_HN(0, 17.265)_ = 0.82, RR_1/3<B<3_ = [1, 49]).^3^

For Jenkins & Obhi (2021), the raw data of 144 participants are available. In the 3D condition at 200 ms, we obtained ΔIB = −0.007022 (slider units) = -7.022 ms with SE = 0.005931 (slider units) = 5.931 ms. The BF analysis, using a half-normal distribution with a standard deviation of 17.265 ms, indicated inconclusive evidence for a stronger binding in near space compared to far space (BF_HN(0, 17.265)_ = 1.06, RR_1/3<B<3_ = [1, 62]).^4^

Together, these direct tests indicate that when the correct ΔIB contrast is applied, neither dataset supports the claim that spatial distance increases intentional binding. Thus, we argue that the available evidence does not suggest that spatial distance modulates temporal bindings in contexts involving intentional movements – i.e., the temporal binding between active movement and passive movement or between active movement and external event baseline tasks. Indeed, no study to date has provided substantial evidence that the difference between the active task and the passive task (Mariano et al., 2024), or the external event baseline task (Jenkins & Obhi, 2021), is modulated by distance. Furthermore, our study also offers inconclusive evidence for an effect of distance on “intentional binding”.

### So, what does spatial distance modulate in these studies?

Both Jenkins and Obhi (2021) and Mariano et al. (2024) reported that the shortest interval was estimated significantly shorter in near space than in far space. In Mariano et al. (2024), this effect was observed in the active movement task, while in Jenkins and Obhi (2021), it occurred in the 200 ms delay, 3D perspective condition, independently of the presence or absence of active movement.

To more directly compare our data with these two previous studies^5^, we conducted a 2 x 3 x 3 repeated-measures ANOVA with the factors Distance (near, far), Task (active movement, passive movement and external sensory event), and Delay (150, 450 and 750 ms). The results involving the Distance factor showed no significant main effect of Distance (F (1, 32) = 2.04, p = 0.16, partial η^2^ = 0.06) and no significant three-way interaction (F (4,128) = 0.57, p = 0.68, partial η^2^ = 0.02). However, the interaction between Distance and Delay (F (2,64) = 3.43, p = 0.038, partial η^2^ = 0.10) and between Distance and Task (F (2,64) = 4.15, p = 0.02, partial η^2^ = 0.12) were significant. Post-hoc analyses revealed that, independent of the task, both the 150 ms (p=0.016) and 450 ms (p=0.028) delays, but not the 750 ms (p=0.31) delay, were perceived as shorter in near space compared to far space. Additionally, independent of delay, the intervals were perceived as shorter in near space compared to far space for the external event baseline task (p=0.048), but not for the active (p = 0.079) and passive (p = 0.073) movement tasks.

Thus, across studies, an effect of proximity seems to emerge: the shortest delay was perceived as shorter in near space than in far space. Such shortening was only observed for the active task in Mariano et al. (2024) and it was also observed, but was independent of the task, in both Jenkins and Obhi (2021) and in our study. In addition, such shortening could also be observed for the middle delay in our work. Importantly, independent of the delay, the near vs. far shortening was also observed in tasks that did not involve any movement, such as the external event baseline task in our data. Therefore, on one hand, the distance-related modulation seems to be consistently observed across studies; on the other hand, it appears to be unspecific to voluntary action execution and not limited to the shortest interval estimation, possibly suggesting that distance may broadly impact the perception of temporal durations.

These findings are also consistent with previous studies using different paradigms. For example, Anelli et al. (2015) employed a duration bisection task and found that durations were judged as shorter in near space. Petrizzo et al. (2021) extended this finding using a duration reproduction task and found systematic under-reproduction of durations in near space and over-reproduction in far space. In contrast, Qi et al. (2019) applied a temporal reproduction task with attentional anchoring to the hands and found that reproduced intervals were longer when stimuli appeared near both hands. This apparent dilation in near space was attributed to prolonged disengagement of attention near the body, especially under sequential response conditions where participants had to respond with both hands; the distance effect was reduced when only one hand was used, supporting the role of attentional anchoring. Finally, spatial modulation of time perception may also impact other processes: for example, Iachini et al. (2017) used a collision prediction task and found lower temporal discrimination accuracy and a more liberal “collision-imminent” bias in near space. Taken together, despite some divergence in the direction of duration perception (i.e., compression or dilation), these studies converge on one point: spatial distance, in and of itself, modulates the perception of time, even in the absence of voluntary action, motor signals, or causal relationships.

Although Mariano et al. measured explicit agency in their study and found no effect of distance, the task used to measure explicit agency was very simple (pressing a button to switch on a lightbulb). Such simple tasks are commonly used in the field, yet they might not be sensitive enough to detect the effect of distance. Moreover, the lightbulb in the “far” condition was not particularly distant, and people are accustomed to this everyday situation, as lightbulbs are usually much farther from the light switch in real life. Therefore, it might be hasty to conclude that spatial distance has no effect on the explicit sense of agency. Therefore, while three studies - Jenkins and Obhi (2021), Mariano et al. (2024) and our own (Kong et al., 2024) - contribute valuable insights into the potential influence (or absence of influence) of spatial distance of an action outcome on the temporal binding and the sense of agency, further investigations are needed to establish whether spatial distance - as manipulated through depth cues, perspective, or VR - impacts the sense of agency.

### Future Directions

To date, three studies, Jenkins and Obhi (2021), Mariano et al. (2024) and our own (Kong et al., 2024) - aimed to examine whether spatial distance of action outcome influences the sense of agency. Despite their methodological differences, all studies pointed to preliminary indications that distance may influence temporal interval perception, irrespective of variations in time intervals or experimental conditions, and this effect does not appear to be linked to agency.

Currently, our lab is actively exploring new experimental paradigms to further investigate the relationship between spatial proximity, the sense of agency, and decision making in voluntary actions (Kong et al., 2025). One key approach, already applied by Mariano et al., involves acquiring both implicit and explicit task paradigms. However, given the increasing challenges that the ‘intentional binding’ paradigm faces as an implicit measure of agency, we would suggest using more natural, complex, goal-directed tasks (e.g., object manipulation, steering) not merely having participants press a button. This will allow for capturing both participants’ conscious and reflective sense of control over their actions and outcomes, and more implicit and automatic sensorimotor performance (Dewey & Knoblich, 2014; Moore et al., 2012; Synofzik et al., 2008). This approach could involve tasks that require continuous control and adaptation (Applebaum et al., 2025), providing a more nuanced understanding of how implicit and explicit aspects of the sense of agency operate in different spatial contexts.

Advancements in VR technology offer exciting opportunities to explore how environmental contexts, whether real or simulated, interact with spatial proximity to influence agency. In VR settings, one can manipulate spatial parameters with greater precision and introduce novel scenarios, such as dynamic environments or interactions with multiple objects (Kong et al., 2017), that are not easily achievable in traditional lab settings. This will allow investigating whether and how spatially complex and immersive environments impact the sense of agency, both implicitly and explicitly. However, such designs can elicit demand-conforming responses from participants. To mitigate these effects, researchers could use a neutral, task-relevant cover story that does not disclose the study’s agency-related aim, brief pre-/post-task expectancy probes with pre-registered criteria for exclusion, and manipulation checks confirming that distance/depth cues are encoded without cueing the hypothesis. Beyond behavior, future research should also examine the neural correlates of spatially modulated agency using neuroimaging techniques, such as EEG, MEG, brain stimulation and fMRI. These methods will help identify the brain processes and neural correlates underlying the putative effects of spatial proximity on agency, providing a more integrated view of how spatial cognition and agency may interact.

Additionally, examining individual differences in the sense of agency would be commendable, as factors such as personality traits, cognitive styles, and motor abilities may influence how spatial proximity affects one’s perception of agency. Understanding these individual differences could have important implications for designing interventions or applications in fields such as rehabilitation, gaming, and human-computer interaction, where the sense of control over distant or virtual objects is crucial.

Finally, it would also be important to investigate whether spatial proximity influences the sense of responsibility over moral action outcomes, as the sense of responsibility is closely linked to agency (Gallagher, 2012; Haggard & Tsakiris, 2009). Exploring whether physical or even psychological distance affects moral responsibility over action consequences could offer insights into how spatial factors might modulate ethical decision-making and accountability in both virtual and real-world settings (Darley & Batson, 1973). This line of inquiry could also build on findings in social psychology that demonstrate the impact of psychological distance on moral judgments, such as construal level theory and its effects on perceived agency and responsibility (Eyal et al., 2008; Trope & Liberman, 2010).

Finally, adversarial collaborations are a promising way to resolve theoretical, empirical and methodological disputes. A Peer Community In Registered Report (PCI RR) ^6^ could preregister: (i) standardised stimulus, hardware and depth-cue calibration; (ii) a Bayesian analysis plan with informed priors plus equivalence testing; and (iii) blinded data processing with code-frozen pipelines. The participating labs could co-develop the Stage-1 protocol, specifying sampling rules a priori.

Together, incorporating these diverse approaches will not only deepen our understanding of whether and how spatial context modulates the sense of agency, but also advance the field by addressing methodological gaps and expanding the scope of investigation to more ecologically valid and technologically advanced settings.

## Data availability

All primary data are publicly available on the Open Science Framework repository (https://osf.io/ua3nw/).

## Acknowledgements

The authors would like to thank the editors and all four reviewers, especially Reviewer 4, for their detailed and helpful suggestions that greatly improved the paper. The authors would also like to thank their team members for the discussion during the team meeting.

## Footnotes

1 Based on values from Supplementary Table S1 of Mariano et al. (2024), the size of the distance effect on temporal interval estimation in the active task was 34.5 ms - twice the size reported in their Figure 2. Re-calculating the BF by using a half-normal distribution with a standard deviation of 34.5 ms, we found substantial evidence that spatial distance did not modulate the temporal binding between active movement and external event tasks (BF_HN(0,34.5)_ = 0.23, RR_B<1/3_ = [23, >3000]). When testing each interval separately, there was also substantial evidence supporting the absence of distance modulation for the 150 ms interval (BF_HN(0, 34.5)_ = 0.28, RR_B<1/3_ = [28, >3000]). For the 450ms and 750 ms delays, the evidence was inconclusive (450ms: BF_HN(0, 34.5)_ = 0.41, RR_1/3<B<3_ = [1, 43]; 750ms: BF_HN(0, 34.5)_ = 0.47, RR_1/3<B<3_ = [1, 50]). In contrast, there was substantial evidence that spatial distance modulated the temporal binding between passive movement and external event tasks (BF_HN(0,34.5)_ = 13.36, RR_B>3_ = [4, 178]), with temporal binding being smaller in near space than in far space – thus opposite to the expected distance effect. When testing each delay separately, there was also substantial evidence supporting the distance modulation in the opposite direction for the 150 ms delay (BF_HN(0, 34.5)_ = 3.93, RR_B>3_ = [5, 47]). For the 450 ms and 750 ms delays, the evidence was inconclusive (450ms: BF_HN(0, 34.5)_ = 0.77, RR_1/3<B<3_ = [1, 86]; 750ms: BF_HN(0, 34.5)_ = 2.46, RR_1/3<B<3_ = [1, 398]).

2 In the Supplementary Table S1, SE_Δ_ = 16.48 ms; as other values were twice the values reported on their Figure 2, we estimated SE_Δ_ = 8.24 ms for the main BF calculation.

3 Using their Supplementary Table S1, we obtained ΔIB = -12.36 ms (i.e., twice the value reported in their Figure 2) with SE_Δ_ = 16.48 ms. Using a half-normal distribution with a standard deviation of 34.5 ms (the size of the distance effect on temporal interval estimation in the active task extracted from their Supplementary Materials, i.e., twice the value reported in their Figure 2), the results was virtually identical: BF_HN(0, 34.5)_ = 0.82, RR_1/3<B<3_ = [1, 99].

4 A BF analysis, using a half-normal distribution with a standard deviation of 34.5 ms (the value indicated in the Supplementary Material of Mariano et al. (2024)) also provided inconclusive evidence for a stronger binding in near than in far space (BF_HN(0, 34.5)_ = 0.59, RR_1/3<B<3_ = [1 62]).

5 To align our summary analysis with the reporting framework used in those studies and to enable direct, like-for-like comparison of omnibus effects and simple contrasts on interval estimates (i.e., outside the intentional binding question), we report a conventional repeated-measures ANOVA in this subsection. Our primary inferential tests for agency claims remain Bayesian and are conducted on the decisive ΔIB contrast in the above section; the ANOVA here serves only to situate distance-related changes in general time-perception.

6 https://rr.peercommunityin.org/PCIRegisteredReports/

